# Agent-based model of multicellular tumor spheroid evolution including cell metabolism

**DOI:** 10.1101/431163

**Authors:** Fabrizio Cleri

## Abstract

Computational models aiming at the spatio-temporal description of cancer evolution are a suitable framework for testing biological hypotheses from experimental data, and generating new ones. Building on our recent work [J Theor Biol 389, 146-158 (2016)] we develop a 3D agent-based model, capable of tracking hundreds of thousands of interacting cells, over time scales ranging from seconds to years. Cell dynamics is driven by a Monte Carlo solver, incorporating partial differential equations to describe chemical pathways and the activation/repression of “genes”, leading to the up- or down-regulation of specific cell markers. Each cell-agent of different kind (stem, cancer, stromal etc.) runs through its cycle, undergoes division, can exit to a dormant, senescent, necrotic state, or apoptosis, according to the inputs from their systemic network. The basic network at this stage describes glucose/oxygen/ATP cycling, and can be readily extended to cancer-cell specific markers. Eventual accumulation of chemical/radiation damage to each cell’s DNA is described by a Markov chain of internal states, and by a damage-repair network, whose evolution is linked to the cell systemic network. Aimed at a direct comparison with experiments of tumorsphere growth from stem cells, the present model will allow to quantitatively study the role of transcription factors involved in the reprogramming and variable radio-resistance of simulated cancer-stem cells, evolving in a realistic computer simulation of a growing multicellular tumorsphere.

## 1 Introduction

In the attempt to model tumor growth *in vitro*, sphere-forming assays, or **tumorspheres**, are a peculiar culture method that allows stem cells to grow in all directions, within a hydrogel scaffold mimicking the natural extracellular matrix structure [1–3]. Compared to ordinary 2D (or “dish”) cultures, 3D-spheroidal cultures generate unique spatial distributions of nutrients and oxygen in the cells, mimicking much better the *in vivo* conditions [4]. Cell lines grown in 3D display different expression profiles, especially for those genes that play a role in proliferation, angiogenesis, migration, invasion, radio/chemosensitivity. Because of their nature and structure, such bio-objects can be (relatively) easily manipulated and characterized by biological techniques, optical microscopy, and also by micro-mechanical tools [5]. Tumor spheroids ideally represent an upper (macroscopic) level of definition of the problem of detecting, following, and quantifying the local and long-range outcomes of stem cell evolution, eventually coupled to DNA chemical/radiation damage. It is worth noting that, while the effects of ionizing radiation have been studied in multicellular tumor spheroids already from the earliest applications of this method in the late’ 70s [6], nothing seems to have been published yet concerning irradiation of scaffold-grown cancer stem-cell spheroids, and very little on tumor-explanted organotypic spheroids [5].

The subject of the present work is the development of a 3D computer simulation model, to be ultimately coupled with the experimental results from stem-cell spheroid cultures. Such a close experiment-theory coupling has been often advocated as a crucial ingredient to inform biological hypothesis making and experimental design [7, 8]. However, the spatial and temporal evolution of a tumor mass, starting from the smallest aggregate of cells (in principle, just one), up to arriving at a macroscopic cancer, is a subject that still largely escapes the possibility of a mathematically-grounded scientific prediction (see e.g. [9, 10]). The morphological growth and development of a tumor results from many factors that are difficult to describe in equations, within a coherent mathematical framework, whether intracellular (such as the adhesion between cells and with the matrix, individual genetics of apoptosis, necrosis, signalling and repair cycles, cell diversity), or extracellular (such as the space-time distribution of oxygen and nutrients, as well as the mechanical stresses on tissues). On the other hand, the availability of mathematical models capable of such a predictive power is very desirable, both on the scale of laboratory research, as well as on the preclinical, and even clinical scale (see e.g. [11–15]). Such models, necessarily intended for the numerical simulation on very large computers, in addition to saving a quantity of material resources, allow to carry out virtual experiments that would be much more difficult, or even impossible to implement in the laboratory.

The various phases of tumor growth have already been the subject of numerous mathematical studies, through the so-called “continuous models” (see e.g. [16–18]), which made it possible to generally formalize some common characteristics, such as: (i) a rapid phase of avascular growth (logistic or Gompertz equation), up to the limit of diffusion; (ii) the phase of angiogenesis, characterized by the diffusion and degradation of the TAF factors, as well as the mobility of the cells attracted by chemo-taxis to the growth region; and (iii) the metastatic phase, characterized (mathematically) by the spatial heterogeneity of cell growth. However, it is practically impossible to take into account the wide variety of conditions encountered by a realistic population of cells within a spatially-continuous model, by its own nature intended for modelling an *average* population of cells. The ambitious objective would be predicting with mathematical rigor the tumor growth in time and space, and the effect of chemo- and radiotherapy treatments, on both cancerous and healthy cells, over time scales ranging from a few hours or days up to long-term follow-up, reaching even years after the treatment.

A practical response may come from discretized biophysical models, in which each cell is described individually with a set of characteristics (descriptors, or “degrees of freedom”) that evolve locally throughout the simulation time, under the influence of internal constraints and stresses coming from local neighbourhood (other cells), or external ones (extracellular matrix, therapeutic treatments, environment) [15, 19]. While obviously more expensive than continuous models in terms of computer resources, a discretized model allows a detail in the description of biological functions, and calibration on experimental data, impossible to obtain by any other simulation model. Moreover, in such a computer model it is relatively affordable to extend the duration of the simulation over very long time intervals, and to study very long-term effects with a flexibility that is rarely accessible to experimental studies. For example, one can repeat the same virtual computer experiment several times, over and over, by changing parameters each time according to the various assumptions formulated by the biologist. The price to pay may be a limited degree of realism, whose quality depends importantly (but not only) on the level of refinement of the initial model calibration phase on real cell lines.

The “agent-based” model (ABM) developed in this work aims at assembling the most relevant biological features in a realistic platform, for the virtual modeling of the long-term evolution of cell proliferation and damage, following various types of “therapy-like” treatments. While some work in this direction has been initiated in the chemo therapy domain [20], it is worth noting that the coupling of agent-based models with ionizing radiation and radiotherapy, to simulate the action of external agents on cancer growth and/or arrest, is not much developed. Even the most recent attempts in this direction (see e.g. [21,22]) did not include explicit simulation of the radiation damage, but rather assumed a pre-existing damage model (such as the linear-quadratic, etc.). An original contribution of the present ABM model (already present in its previous 2D version [24]) is therefore the inclusion of explicit damage accumulation and repair, at the single-cell level, coupled to mathematical evolution of expression/repression markers in the cell cycle.

Spatio-temporal modeling of chemical concentration evolution by partial differential. equations (PDEs) is one of the established techniques in the field of cancer and tumor growth modeling (see e.g. [23] and references therein). The cell/agents of our extended ABM include intra-cell metabolic networks described with this more classical technique, and the resulting space-time coupled (”reaction-diffusion”) equations are solved numerically; cell-to-cell communication is based on exchange of input and output quantities issued from such intra-cell PDE networks, which accumulate in the respective source terms.

## 2 Agent-based model of cell evolution

The 3D-ABM we are developing with the in-house code MODLOG [24], already allows to consider many important applications in the context of the simulation of the tumor evolution, as well as the study of the effects of radiation and chemical treatments on the whole of the cell population (healthy, cancer, stem cells). The virtual cells (”agents”) of the theoretical model have the possibility to change their state following localized events, to exit in a state of dormancy in which their activity is stopped, although their presence continues to influence the evolution of the neighbouring population, or undergo transformation, neoplastic and apoptosis.

The first published version of our ABM [24] allowed a direct comparison with experimental results of 2D culture growth, with a 1-to-1 correspondence both in time and length scales. Real cell transformation events are thought to follow a Poisson process [25], therefore it is justified to model both cell evolution and induced damage as a Markov chain [26, 27]. We assume that agent-cells can be in any state *n∈* [0*, m*], with *n*=0 corresponding to a pristine cell with zero accumulated damage, and *n*=*m* to a cell with a maximum of accumulated damage. The time-dependent behaviour of each cell *c ∈N* from a population of *N* agents is characterised by a number of descriptors (phenotypes), collected in a state vector:

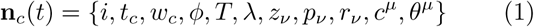

for the *ν* = 1*,…, k* different types of lesions (e.g., single and double strand break, mutation, abasic site, etc.); *i* is the lattice site occupied at time *t*; *t_c_* is the local clock of each cell; *w_c_* the fictitious cell volume; *ϕ* is the cell phase, running through G1, S, G2 and M; *T* is the cell type (epithelial, fibroblast, cancer, stem, etc.); *λ*=1,2,…7 is the cell state, the increasing index values respectively indicating a normal, senescent, quiescent (or G0), arrested, neoplastic, stem, or dead cell; *z_ν_* is the number of accumulated damages of type *ν*; *p_ν_* is the corresponding damage probability; *r_ν_* the repair probability; *c^μ^* and *θ^μ^* the concentration, and diffusion time, of each chemical species *μ*. The reader is referred to our previous work, for the full details of the basic mathematical features already included in the model [24].

In the present 3D development, the simulation space is filled by a continuous network of Voronoi polyhedra (VP, Figure 1). A VP can be empty, or occupied by a cell. The VP centers are generated by means of Bridson’s algorithm [28], by placing points at random in a right-prism or spherical-shaped volume; starting from the volume center, each new point is added within a fixed radius *R* from the previous ones, with the condition of not being closer than 2*R* to any of the previously placed points. In this way, the simulation space can be densely filled by a random, but homogeneous ensemble of points. Subsequently, the Delaunay tessellation around the set of points is obtained, and the dual set of VPs is constructed. This construction is completed only for the points lying within a non-periodic volume inscribed within a concentric portion of about 2/3 of the whole simulation space. In this way, we can make sure that all the Voronoi polyhedra contained in this sphere have a tightly controlled average size, and average connectivity to a mean number of nearest neighbors. In practice, for an average cell size 2*R* ≃ 15 *μ*m, about 3,5 million cells can be closely packed in a cubic simulation volume of ~2,000^3^ *μ*m^3^, corresponding to a maximum enveloping spherical volume of about 1 mm radius. This size is largely sufficient to simulate the time evolution of realistic neurospheres, which typically contained less than a few million cells in the earlier experiments [29, 30], while are restricted to a few hundreds or thousands units in modern experiments on stem cells [1, 3].

**Fig. 1.**
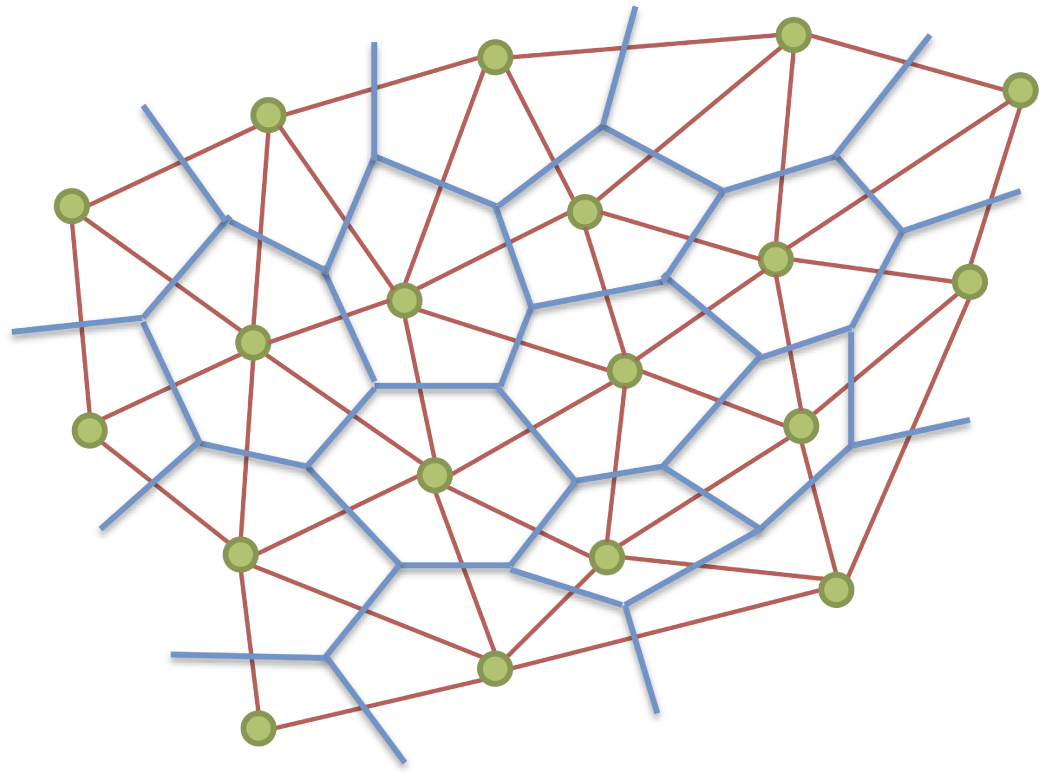
Scheme of the 3-dimensional cell simulation space filled by Voronoi polyhedra. 2D projection of the 3D development on a fixed random grid of points generated by a Bridson algorithm (green dots), which represent the vertices of a Delaunay triangular tessellation (red thin lines). The centers of the Delaunay triangles are the vertices of Voronoi polyhedra (blue lines), these latter representing either filled or empty cell voxels.

The ensemble of VP represents the “world” in which cells live, can multiplicate, evolve, and die. Empty VPs make up the extracellular matrix. We took the approximation of a VP structure that does not change over time, but rather represents a kind of fixed random lattice, in which a cell can eventually move by jumping between neighbor VPs. Such an approximation saves a large amount of computing time, while having a minor impact on the sphere evolution (it may be noted that this same approximation would be entirely inappropriate for, e.g., morphogenesis modelling). During migration, cells interact with all their neighbors in direct contact, and even with remote sites by the long-range diffusion of various chemical species. The important notation is that cell properties are not bound to the local site they are temporarily occupying, but follow (i.e., are carried around) by each cell during its displacements. Notably, one of the parameters in the cell state vector (see above) corresponds to the fictitious volume attributed to each given cell; subject to proper constraints, this allows small changes in the cell-cell distance.

### 2.1 Coupled glucose/oxygen system network

Mathematical modeling of metabolism at the scale of the single cell is a widely developed subject, which gained in complexity in parallel to the increase of computing power [31, 32]. Molecules and pathways involved in signal transduction have been identified and their function understood. Identification and analysis of protein network motifs led to understanding of how molecular interactions function to suppress noise, amplify signals, or provide robustness (see e.g. [33] and references therein).

By comparison, the level of communication among cells is rather poorly understood [34]. Inter-cell communication networks process input signals through intra-cell networks, to obtain an output representing a change in the state of each cell, as well as an input signal to other cells. In the words of Thurley et al., “cell-to-cell communication networks are *networks of networks*”, with many different cell types. Whereas the well-known rules of chemical kinetics apply to the intra-cellular building blocks, it is still unclear how best to model cell-to-cell communication networks.

In developing our multi-cellular model, we aim at the objectives of: (i) establishing a basic connection between oxygen and nutrients (glucose) consumption by the cells; (ii) linking the evolution of these metabolites to the energetic cell cycles, and notably to the ATP/ADP ratios measured in normal vs. cancer cells; (iii) linking these basic networks to other potential cancer markers, in order to build computer simulations of multicellular growth comparable to experimental tumorsphere data. In the present work we focus on items (i) and (ii), leaving the vastly complex (iii) to forthcoming works. In this Section 2, we describe two mathematical models: an “extended” model, in which cell-level glycolisis, carboxylic acids and respiratory cycles are accounted; and a “reduced” model, in which most of the intermediate steps of the energetic cycles are lumped into symbolic reaction paths, with only oxygen, glucose, ATP and ADP being explicitly tracked. The two models are, respectively, more complex and less complex than the Casciari-Sotirchos-Sutherland model [29], which is usually considered a reference in zero-dimensional modeling of neurospheres. In particular, the “reduced” model does not explicitly track the pH of the cell, which may be a considerable over-simplification in some problems. For the sake of computational efficiency only the second, simplified model is the one actually implemented in the ABM applications that will be presented in Section 3 below.

In the ABM, each cell is fed with nutrients and oxygen from the exterior matrix. A coupled system network is set up at the single cell scale, to define the individual viability condition for each cell. The variables in the cell network are not meant to faithfully represent each individual chemical component, but rather the main steps in the chemical reaction paths, in a synthetic way. A graphic summary of such a coupled glucose/oxygen system is represented in Fig 2. The consumed species are glucose [G] and molecular oxygen [O], which enter the system by diffusion with the respective coefficients *D_G_* and *D_O2_*.

**Fig.2.**
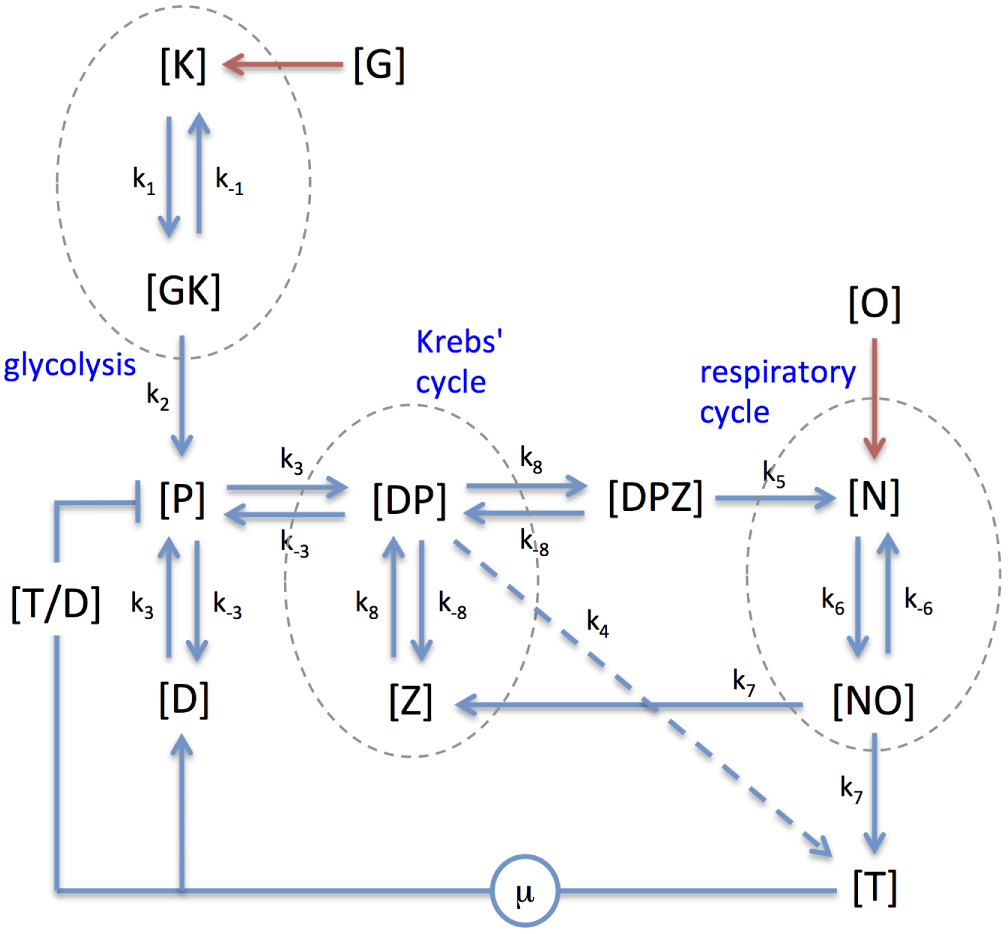
” Extended” model of the coupled glucose/oxygen system network. One-letter symbols such as [X] are the concentrations of individual species. Two- and three-letter symbols such as [XY] indicate a bound complex. Red arrows indicate the input of consumed species. Blue arrows indicate enzymatic reactions; the dashed-blue arrow indicates anaerobic pathway; the ‘⊣’ symbol indicates a repressor feedback. The term *μ* represent fuel (ATP) consumption. (Color online)

In this “extended” model of cell respiration, either completely aerobic (full lines in the figure), or anaerobic (dashed) pathways can be taken by each cell, according to its local conditions and food/oxygen supply. Our starting system of 11 time-evolution equations is:

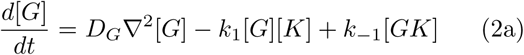

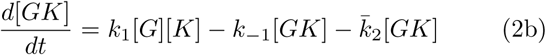

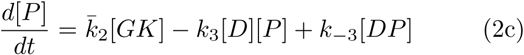

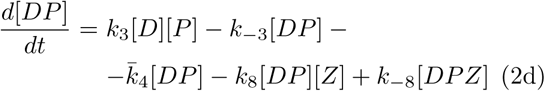

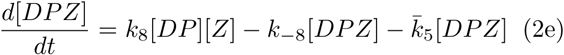

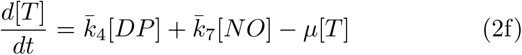

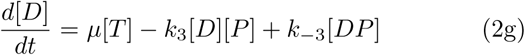

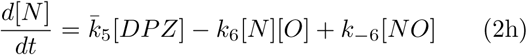

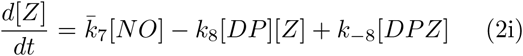

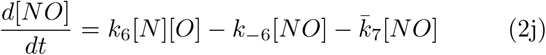

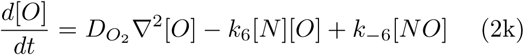

The *K_m_/K_–m_* indicate the rates of reversible reactions, whereas irreversible ones are indicated with an overbar, *k*̄_n_. The (2a) represents input of glucose minus a fraction activated by PFK protein [K], plus a contribution from the dissociation of the [GK] complex. The (2b) is formation/ dissociation of [GK] minus a fraction (proportional to the concentration, by law of mass-action) going into the end-product [P] (pyruvate). The (2c) represents [P] making a complex with ADP,

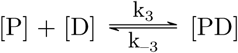

while being supplied by the previous step (2b). The two steps (2a)-(2b) make up the **glycolysis**.

The (2d) describes binding of the [PD] complex to NAD+ or FAD^2+^(both indicated by [Z]),

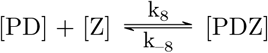

while a fraction of the complex gives off ATP at a rate k̄_4_ (for the sake of simplicity, this effectively cumulates also the ATP produced in glycolysis). The (2h) describes the output of NADH (indicated by [N]) at a rate *k*̄_5_, for simplicity including also FADH_2_ reduced from FAD^2+^). This ends the **Krebs’ cycle**.

In the subsequent aerobic cycle, [N] combines with molecular oxygen [O], Eq. (2j), supplied by diffusion (2k), at a rate *k*_6_ (and dissociates at a rate *k_−_*_6_). The as-formed [NO] complex is consumed in the **respiration**, Eq. (2h), and gives off ATP at a rate *k*̄_7_, while the reduced co-enzymes are oxidised back to NAD^+^/FAD^2+^. If the anaerobic (dashed) pathway is selected, considerably less ATP is produced at a rate *k*̄_4_ directly from the [DP] complex, and no NAD/FAD are cycled. Either way, ATP is consumed by cellular processes (see below) at some rate *μ*, to be individually specified during the 3D cell evolution, and turns back to ADP.

Since [K] is explicitly treated as an enzyme, no time evolution equation is written correspondingly, however its concentration is governed by the conservation condition:

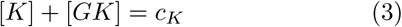

allowing to rewrite Eqs.(2a)-(2b) as:

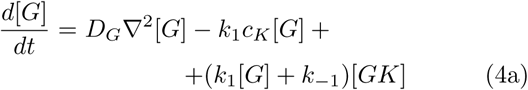

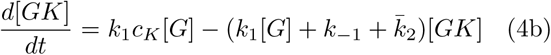

The rate of glycolysis is back-regulated by the ratio ATP/ADP (see the [T/D] repressor in the figure): a high concentration of ATP reduces the glycolysis rate; conversely, when the ATP concentration falls, an increase in glycolysis is activated. In practice, the [T/D] ratio governs the value of the rate constant k̄_2_ as:

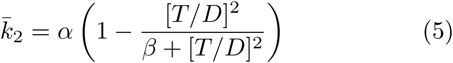

with *α, β* adjustable parameters.

The tests carried out with this extended model gave rather complex results, since the PDE system is quite unstable; the range of parameters for which a physically motivated result appears is extremely narrow, and sensitive to the smallest variations. However, our main objective here is to control the glucose and oxygen consumption in each cell, by linking it to the demand for ATP. Therefore, many of the intermediate steps may become irrelevant, at least to a coarse level of approximation.

### 2.2 Restricted model of glucose/oxygen metabolism

To simplify the complex network of coupled glycolytic, citric acid and respiratory cycles, we drastically reduced the steps to only **two** fictitious intermediates, namely: a bound state, labelled [GD], of [G] (glucose) and [D] (ADP); and a bound state of [GD] and [O] (oxygen); moreover, all the reactions are taken as irreversible (Figure 3). The PDE system is thus reduced to only 5 time-evolution reaction-diffusion equations, as:

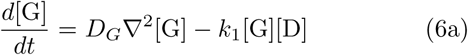

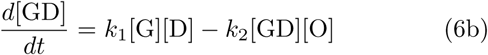

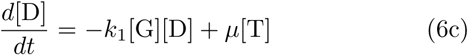

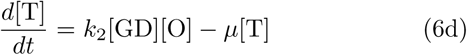

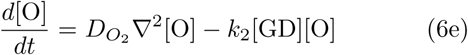

The rate of glucose consumption is back-regulated, between 0 and a maximum *α*, by the ATP/ADP ratio ([T]/[D]) as

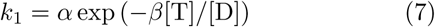

with *α, β* adjustable parameters. To a first approximation, the conversion efficiency of ADP to ATP in the last step is linearly related to the ADP concentration, as:

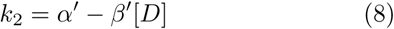

**Fig. 3.**
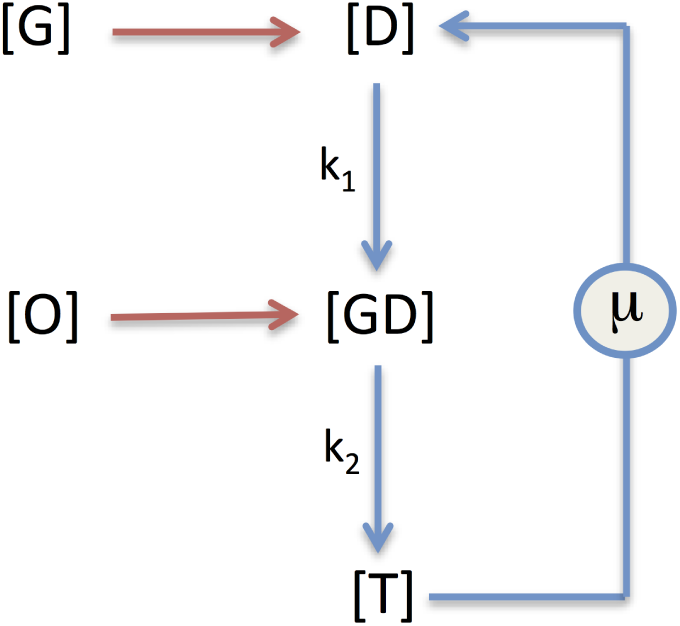
”Reduced” model of the coupled glucose/oxygen system network. See Fig. 2 for the meaning of symbols. (Color online)

The key quantity we are interested in controlling at this stage is the ATP/ADP ratio, under different conditions of oxygen and nutrients supply, as well as with different conversion rates *k*_1_*, k*_2_, and variable ATP utilization rate *μ*. The ATP/ADP ratio is one of the key parameters in identifying the response of healthy vs. cancerous cells [35, 36], the latter being often characterized by a reduced mitochondrial metabolism and lower ATP/ADP ratio that favors enhanced glycolysis.

The simplest model situation is that of a steady-state supply of glucose and oxygen, i.e. *d*[G]*/dt* = *d*[O]*/dt* = 0, for which the condition:

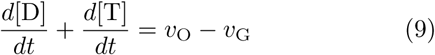

is easily obtained, with *v*_G_ and *v*_O_ respectively the diffusion terms in Eqs. (6a) and (6e); moreover, for vanishing extra-cellular gradients the above condition becomes *d*[D]*/dt* = − *d*[T]*/dt*. Symmetric variations in ATP and ADP could also imposed by adjusting the reaction rate *k*_2_ as:

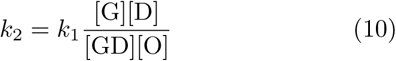

a condition that automatically maintains constant the total [T]+[D] concentration, reducing the system to only four independent equations. Notably, the two rates *k*_1_*, k*_2_, with they back-regulation of glucose and oxygen implicitly maintain the electrolyte equilibrium (lactate / pyruvate, 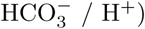. However, explicit tracking of the cell pH is not possible in this approximation.

More interesting situations are obtained by imposing a periodic supply of nutrients, which however requires a numerical solution because of the strong non-linearity of the system of equations. By numerical testing, it was possible to identify domains of stability of the system (6a)-(6e) for which periodic solutions remain stable over sufficiently long intervals of time. The time step for the explicit integration of the PDE system is 0.1 to 1 minutes, the stability being verified over time intervals of the order of ~10^5^ min. The following results are presented for time intervals of 2 ×10^4^ min, corresponding to about two weeks, a typical time for cell sphere growth experiments.

In Figure 4 we report examples for two different test cases studied, for periodic glucose supply with time profile, respectively, sinusoidal:

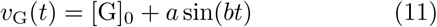

and impulsive with a periodically-defined exponential relaxation time, *t*_0_ = *mτ*:

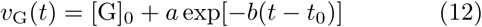

**Fig. 4.**
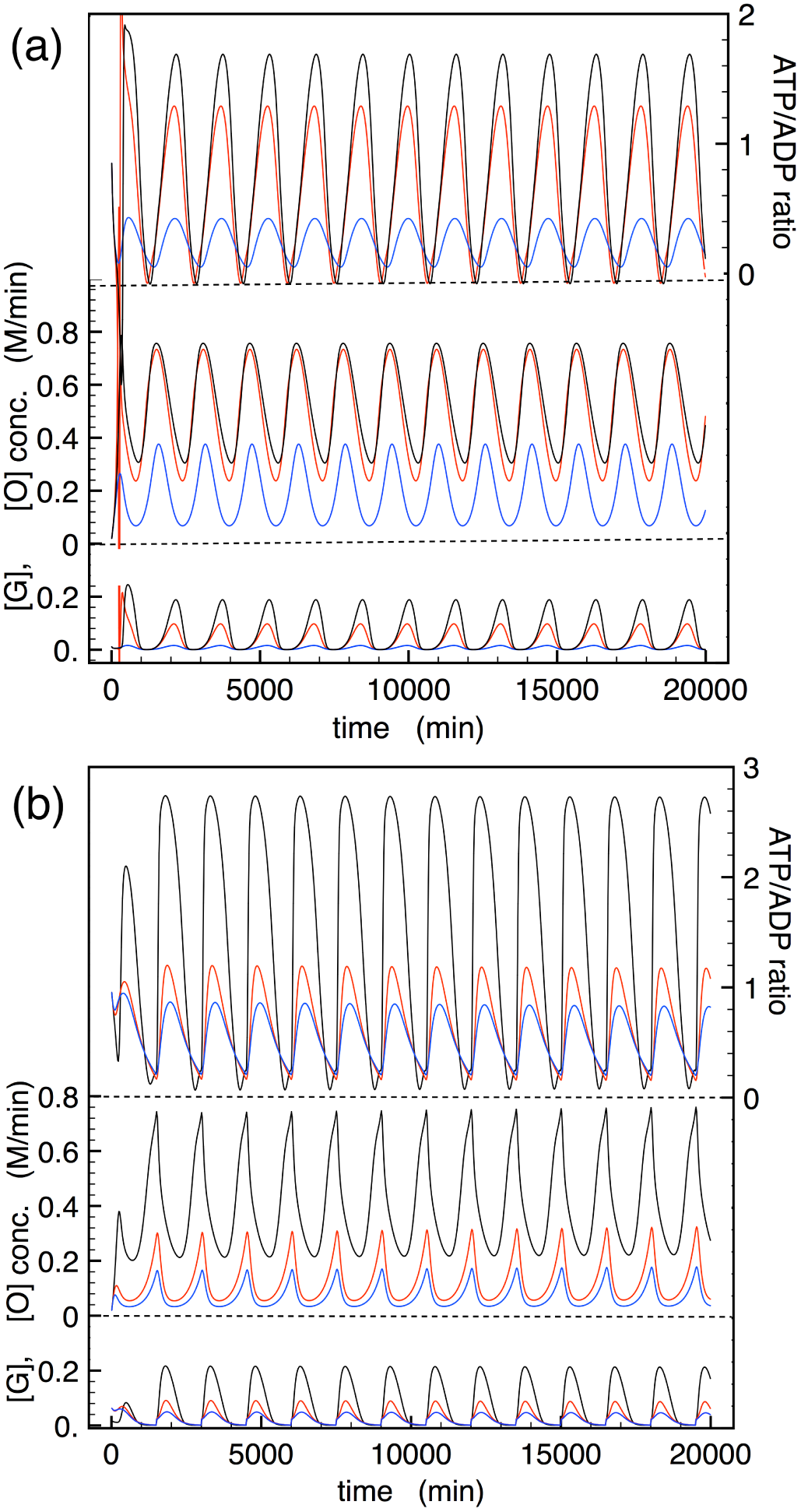
Time evolution of the “reduced” glucose metabolic model. (a) Example of the model with sinusoidal glucose supply. [G] and [O] concentrations are reported on the left ordinates, the ATP/ADP ratio on the right ordinate axis. Black, red and blue curves correspond to (*α^′^, β^′^*)=(0.009,0.04), (0.018,0.08), (0.036,0.08) in Eq.8. (b) Model with impulse supply of glucose, followed by exponential decay. Black, red and blue curves correspond to *α^′^*=0.018, 0.052, 0.078, *β^′^*=0.08, in Eq.8. (Color online)

The latter scheme may be representative of a periodic addition of a fixed amount nutrients to the culture, e.g. on *τ* =24h basis. Table 1 provides ranges of parameters for which stable solutions can be obtained. For all the examples reported we imposed a constant oxygen supply, *v*_O_(*t*)=1 mM/min, and a constant ATP utilization rate *μ*=1 mM/min. We used the initial concentration values (in molar units) [G]_0_=0.01, [O]_0_=0.01, [GD]_0_=0.01, [T]_0_=0.5,[D]_0_=0.5; given the non-linearity of the system, the dependence on the choice of initial values is non negligible.

**Table 1.** Parameter ranges of the model (6a)-(6e) giving the stable solutions of Fig. 4 and 5. Oxygen supply is constant for all variants, at *v*_O_(*t*)=1 mM/min. Note that these ranges would change for different initial values of the baseline model parameters.

| [G] profile | $\mu$ | $a$ | $b$ | $\alpha$ | $\beta$ | $\alpha'$ | $\beta'$ |
| --- | --- | --- | --- | --- | --- | --- | --- |
| sinusoid | [0.01 – 0.05] | [0.001 – 0.008] | 0.004 | [0.1 – 5] | [0.03 – 3] | [0.01 – 0.2] | [0.02 – 0.08] |
| pulse + exp | [0.01 – 0.05] | 0.00315 | 0.002 | [0.1 – 5] | [0.03 – 3] | [0.01 – 0.2] | [0.02 – 0.08] |

The sinusoidal supply of glucose is a condition that gives stable solutions for a rather large interval of the model parameters. Note that the oxygen concentration [O] becomes periodically dependent on time, although its input is strictly constant in these examples. The periodic solutions show relatively little dependence on the rate *k*_1_, while the response in terms of the ATP/ADP ratio is almost entirely contained in the variation of the rate *k*_2_. The curves in Fig. 4a show results for the ATP/ADP ratio, [G] and [O] concentration, for different values of the parameter *a* (for the other model parameters, see Table 1). A parallel increase in the baseline level values of *v*_O_ and *v*_G_ leads to increased ATP/ADP ratio.

From the curves reported in Fig. 4a it can be seen that the [G] concentration at equilibrium is somewhat distorted, compared to the input sinusoidal profile. This modified response is more evident for the lowest ATP utilization rates, *μ~* 1 mM/min, while the profile remains closer to sinusoidal for higher rates, *μ* ∼ 50 mM/min. The PDE system of the “reduced” model is quite stable against variations in the ATP utilization rate. Figure 5 shows a stable solution obtained with a random value of *μ* in the interval [1-40] mM/min, fluctuating with a frequency of 1 min^*−*1^. The ATP/ADP ratio follows the glucose supply rate in a stable periodic cycle, with random fluctuations superimposed.

**Fig. 5.**
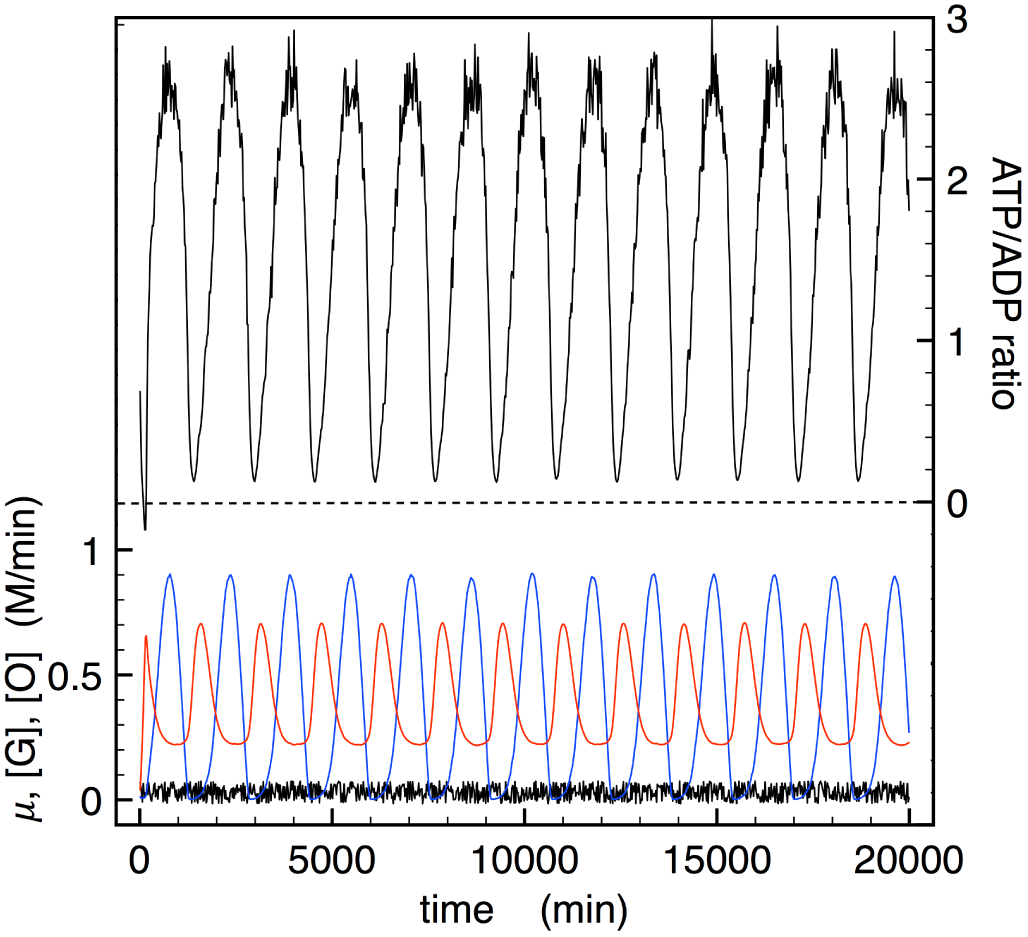
”Reduced” metabolic model with random ATP utilization rate. Glucose supply is sinusoidal and oxygen rate is constant. The ATP utilization rate *μ* (black trace), [G] (blue) and [O] (red) concentrations, are reported on the left ordinates; the ATP/ADP ratio on the right ordinate axis. Simulation parameters: *α^′^*=0.018, *β^′^*=0.06, in Eq.8, *v*_O_(*t*)=3 mM/min, *a*=3 mM/min, *μ* randomly variable in the interval [0.001-0.04] with frequency 1 min^*−*1^. (Color online)

The case with an impulse supply of glucose followed by exponential relaxation, shown in Fig. 4b with examples of stable solutions spanning ATP/ADP ratios from below to well above 1, demands a more subtle definition of the parameters. As for the sinusoidal case, we observe that the [G] concentration at equilibrium differs from the input signal, in that the profile becomes more “rounded”, while retaining the alternance of impulse and relaxation. In this case, however, we often observed a tendency to drift of the oxygen concentration, which tends to steadily increase or decrease with time; only for a narrow window of values of the parameter *a* we could obtain stable solutions, for the chosen values of *v*_O_ and *μ*. Within this parameter window, we again observed that the key parameter in adjusting the ATP/ADP ratio was *α^t^*, the decrease in the ratio corresponding to the progressive increase in the average value of the conversion rate *k*_2_.

## 3 Results of 3D simulations of cell spheroid growth

A tumorsphere is a solid, spherical formation developed from the proliferation of one cancer stem/progenitor cell. Tumorspheres are easily distinguishable from single or aggregated cells [2], as the cells appear to become fused together and individual cells cannot be identified. This assay can be used for example to estimate the percentage of cancer stem/progenitor cells present in a population of tumor cells. The size, which can vary from less than 50 to 250 micrometers, and the number of tumorspheres formed in an experiment, are quantities that can be used to characterize the cancer stem/progenitor cell population within a population of *in vitro* cultured cancer cells, as well within *in vivo* tumors [37, 38]. This method also provides a reliable platform for screening potential anti-CSC agents. The *in vitro* anti-proliferation activity of potential agents selected from tumorsphere assay is more translatable into *in vivo* anti-tumorigenic activity compared with general monolayer culture. Tumorsphere assay can also measure the outcome of clinical trials for potential anti-cancer agents [39]. In addition, tumorsphere assay may be a promising strategy in the innovation of future cancer therapeutics, and may help in the screening of anti-cancer small-molecule chemicals [40].

In this Section we present examples of the 3D-growth simulations of multicellular spheroids with the ABM including local cell metabolism. The simulation typically starts from a small aggregate of a few 100 or 1000 cells, occupying the sites of a Voronoi polyhedra (VP) network as described above. Cells are aggregated in a spheroidal or prismatic starting mass, surrounded by a large space of empty VP, representing the extracellular matrix (ECM). Initial values of metabolite concentrations ([G], [O], [ATP], [ADP]) are provided in each occupied VP, either homogeneously or randomly distributed. At the start of the simulation, the extracellular matrix provides a external inflow of [G] and [O], the diffusion between ECM and cell membrane being considered instantaneous compared to cell-cell diffusion times. In this way, only the outer cells directly in contact with the ECM receive the initial metabolite input, which is passed down to the inner spheroid volume only by diffusion. (Glucose facilitated diffusion is assimilated to spontaneous diffusion; other modes of active transport will be subsequently introduced in the model, for channel-regulated input and output of more complex metabolites.)

In practice, the Laplacian term in Eq. (6a) and (6e) is calculated numerically, by accumulating the concentration differences between each cell and its neighbors in immediate contact, weighted by a coefficient proportional to the distance between each pair of cells (see [24] for details). Diffusion coefficients are given in units of *δ* = *a*^2^*/τ*, a reduced unit based on the average cell-cell distance in the VP lattice *a*=15 *μ*m, and the (adjustable) cell duplication timescale *τ*; for a typical value of *τ* =100 min^*−*1^, it is *δ*=3.75 *×*10^*−*6^ cm^2^/s. Note that such diffusion coefficients do not refer just to the permeability of the cell membrane, but to some effective combination of cell properties and cell density, which overall determine the ability of nutrients to penetrate the volume of interest.

### 3.1 Simplified tests on diffusion-driven growth

In the foregoing examples, we describe generic cells, with 24h cycle and mitosis allowed in the last ~1h of the cycle. Cells are not synchronized at start, i.e. a random value of the starting phase (G1, S, G2, M) is assigned to each cell. For mitosis to occur, we impose that its [G] and [O] must be above prescribed threshold levels; eventually, the more strict condition that the cell must have adjacent free space (non-confluent), may be added. For a test of the general response of the 3D cell growth model these first simulation examples will not use the full metabolic network, but only glucose/oxygen supply from the ECM, followed by ordinary diffusion, with fixed consumption rates equal for all cells. Constant or periodic glucose supply is simulated, while oxygen supply is always kept constant in the outer matrix.

Different types of tumors are found to display quite variable growth-laws and growth rates, ranging from exponential, to linear, to sigmoidal (Gompertz) with saturation, to power-law [41]. In our study we monitor the time dependence of the spheroid radius as a function of basic parameters. We find that one key parameter influencing the growth mode is the threshold for glucose viability, that is the value of concentration [G]_*V*_ above or below which a cell can, or cannot, duplicate. For a given concentration [G]_0_ in the ECM, it was observed that [G]_*V*_ ≲0.7[G]_0_ results in nearly all cells in the mass proliferating, leading to exponential volume growth; increasing [G]_*V*_ to ~ 0.9[G]_0_ results in a linear growth of the sphere radius *R*(*t*), an obvious effect of limiting proliferation only to cells lying on the outer surface of the spheroid, in contact with the ECM (same results are obtained with the non-confluence condition); and restricting the proliferation to just a few cells on the surface, [G]_*V*_ > 0.95[G]_0_, leads to “atypical” power-law growth of the sphere volume.

In Figure 6 we report the radial profiles of [G] and [O] concentration for a simulation starting from *N*_0_=10,000 cells, with initial spheroid radius *R*_0_=168 *μ*m. Glucose is supplied at baseline value [G]_0_=16.5 mM, instantaneously at time *t*=0 of each day, and its concentration in the ECM follows an exponential decay, going to 1/*e* in 500 min; oxygen in the ECM is constantly maintained at the baseline value [O]_0_=3 mM (see e.g. [30]). The simulation time extends for about six days, and the profiles in the Figure are taken at *t*=0 and 12h of each day. It can be noticed the alternance of glucose concentration, between times *t*=0 (when new glucose is supplied) and 12h. The [O] concentration in the cells at the outer surface of the spheroid remains approximately constantly around the baseline value; the [G] concentration is slightly lower than the baseline value for the cells at the surface, and has its maximum a few *μ*m below surface.

**Fig. 6.**
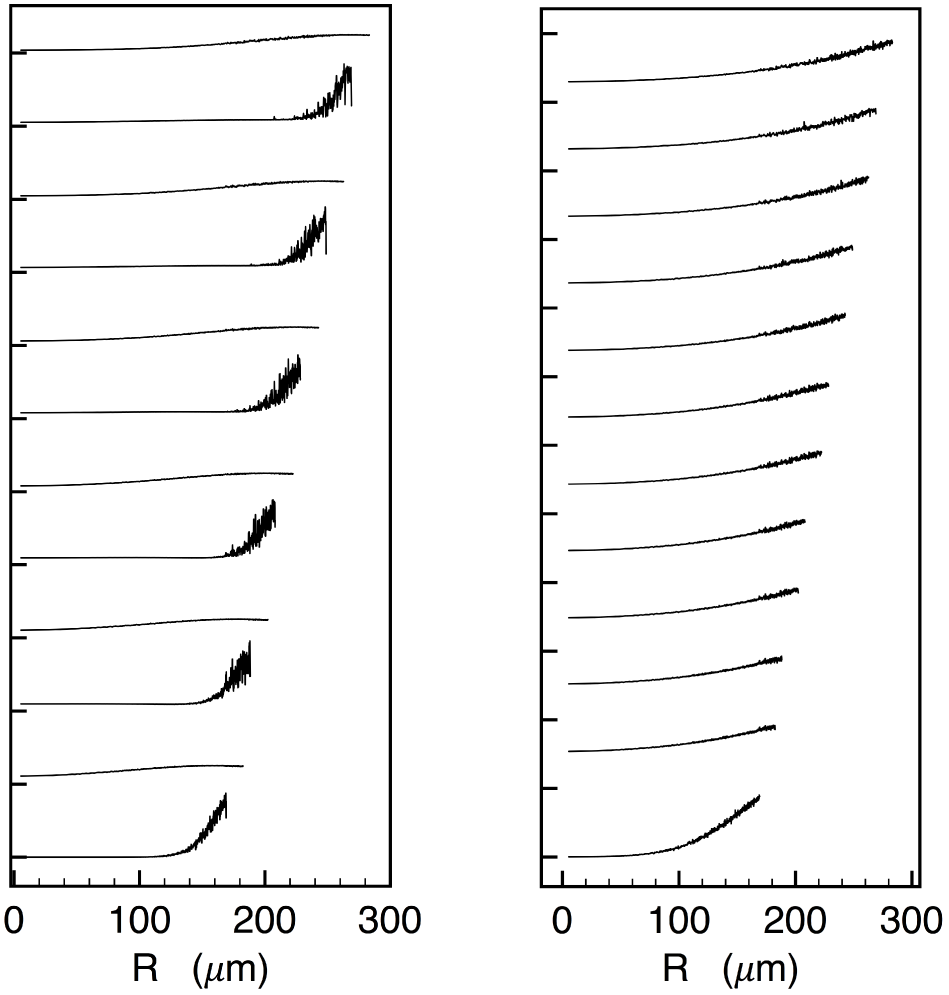
”Reduced” metabolic model with random ATP utilization rate. Glucose supply is sinusoidal and oxygen rate is constant. The ATP utilization rate *μ* (black trace), [G] (blue) and [O] (red) concentrations, are reported on the left ordinates; the ATP/ADP ratio on the right ordinate axis. Simulation parameters: *α^′^*=0.018, *β^′^*=0.06, in Eq.8, *v*_O_(*t*)=3 mM/min, *a*=3 mM/min, *μ* randomly variable in the interval [0.001-0.04] with frequency 1 min^*−*1^. (Color online)

Hypoxia is a major hallmark of cancer cells, increasing the radioresistance compared to well oxygenated tissues. Real tumors have a heterogeneous oxygen distribution, and the same is true for artificial tumorspheres that usually include a viable (oxygenated) outer rim, an hypoxic concentric region, and a central anoxic, necrotic core [42]. Therefore, an important quantity to monitor also in the model is the oxygen radial profile. The dependence of the oxygen profile on the value of the diffusion constant is shown in Figure 7, for *D_O_*2 =0.001, 0.01, 0.1 in reduced *δ* units. This simulation involves only a constant supply of oxygen on the outer surface of a spheroidal aggregate of cells, therefore the concentration profile could be computed analytically for an average cell representative of the growing mass, neglecting the correlations arising in the dynamic effect of the multicellular spheroid growth. Indeed, the results of Fig. 7 closely follow the known laws of purely diffusive behavior (see e.g. Fig. 6.1 of Ref. [43], describing the inward diffusion from the surface of a sphere), which represent a good check of the basic model.

**Fig. 7.**
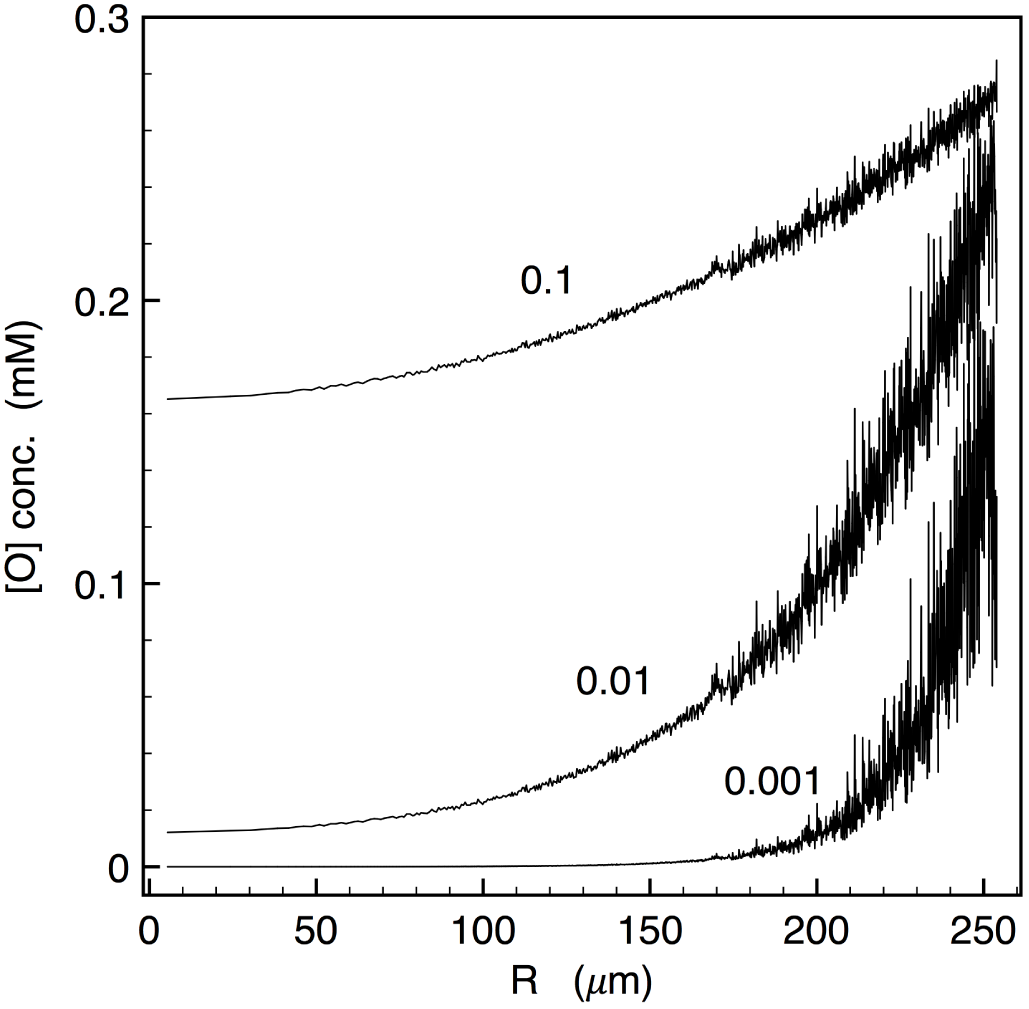
Oxygen concentration profiles. Equilibrium radial distribution of oxygen concentration in a 250 *μ*m cell spheroid. for different values of the oxygen diffusion constant (expressed in reduced units of 3.75*×*10^*−*8^ cm^2^/s).

### 3.2 Tests including the reduced model of metabolism

In this batch of simulations, each cell will also include the “reduced” model of correlated [G] and [O] consumption, resulting in variable [ATP]/[ADP] ratios. Such a signal is interesting per se, in that it allows to check the self-regulation of the system; moreover, it could be further linked to cell viability, e.g. by introducing a “necrosis” factor depending on the ATP availability, and in subsequent developments to a number of other cascade signals.

The next two examples use the same parametrization of the metabolic model, with the following values: *α*=1, *β*=1, *α^′^*=0.04, *β^′^*=0.06; *a* and *b* are not used, since the glucose and oxygen input rates for each cell are dynamically given by the diffusion terms coming from the neighboring cells; the diffusion coefficients are respectively fixed at *D_G_*=*δ* and *D_O_*2 =*δ/*25. The ATP utilization rate is randomly sampled in the interval *μ* ∈ [0.001 0.03] min^*−*1^. In both cases, the oxygen baseline concentration is [O]_0_=0.6 mM. The two examples differ only in the glucose baseline concentration, which is taken to be just slightly above the glucose viability threshold in the first one, and as large as 5 times the threshold in the second one. For a typical threshold [G]_*V*_ =5 mM [30], we set [G]_0_=5.7 mM in the first case, and [G]_0_=25 mM in the second. This difference is enough to give rise to definitely distinct growth patterns.

Figure 8 reports the results for the first run, with [G]_0_=5.7 mM. In this case, only a few cells at the surface will be at [G] values above the proliferation threshold, therefore the growth pattern is surface-limited with linear radius growth rate, *R*(*t*) ∝*t*. Fig. 8a shows that, after an initial stasis, followed by rapid, nearly-exponential rise, from *t ~* 8*d* the spheroid radius starts to increase linearly, as does the cube root of the cell number (*N* (*t*))^1*/*3^. Fig. 8b compares the [G] concentration profiles at *t*=6*d* (red trace), that is in the middle of the rapid growth, and at *t*=10*d* (black trace), that is well into the linear growth regime. It can be seen that the “viable rim” of the spheroid, i.e. the fraction of the spheroid in which cells can proliferate with [G]*>*[G]_*V*_, remains around 0.8*R_max_*; however, the increasingly larger *R_max_* implies that the actual rim thickness is increasing over time. Fig. 8c shows the same data for the [O] concentration; it may be noted that the profile is not purely diffusive, and displays a tendency to approach towards the baseline value in the rim region, for increasing spheroid size. The [ATP]/[ADP] ratio instead, reported in Fig. 8d, does not show appreciable differences during the entire growth time, remaining constantly above 1 with just a little overshoot at the spheroid surface.

**Fig. 8.**
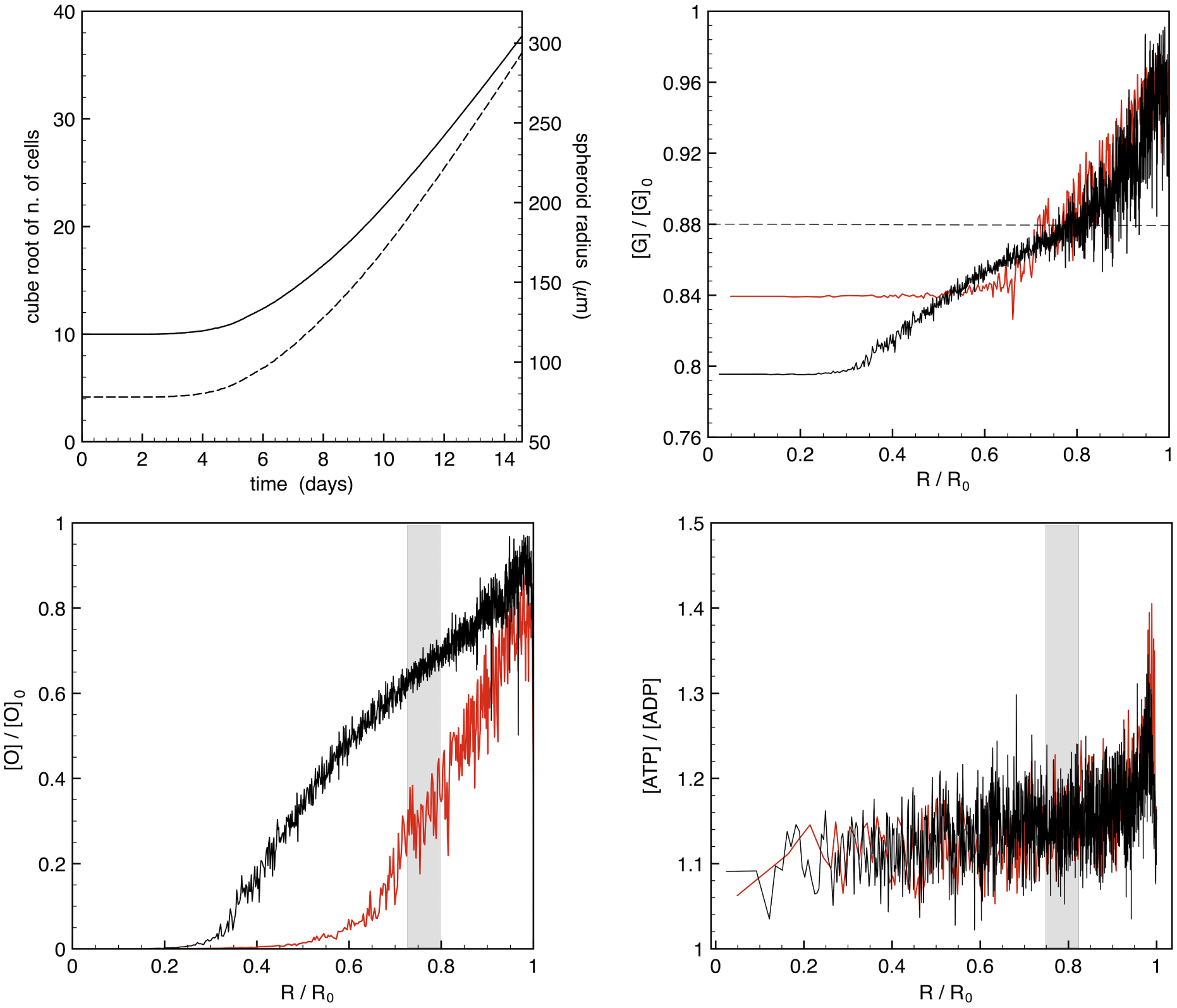
Simulation of 3D spheroid growth in the surface-controlled regime. (a) Growth rate of the cell number (full, left ordinates) and of the spheroid radius (dashed, right ordinates), as a function of time. The linear regime starts around *t* ≃8 days. (b) Glucose concentration ratio (units of the baseline concentration [G]_0_=5.7 mM) at times *t*=6 days (red) and 10 days (black), as a function of spheroid radius (normalized to the respective *R_max_* for each data set). The horizontal dashed line indicates the glucose viability threshold, [G]_*V*_, in this case set at 0.88 [G]_0_. (c) Same as (b), for the oxygen ratio (with [O]_0_=0.6 mM). The vertical grey band indicates the range of *R* values at which [G] cross the value [G]_*V*_. (d) Same as (b), for the [ATP]/[ADP] ratio. (Color online)

Figure 9 reports the results for the second run, with [G]_0_=25 mM. In this case, most cells in the system will be constantly at [G] values well above the proliferation threshold [G]_*V*_. The exponential growth regime is clearly demonstrated in Fig. 9a, in which the cell number is linear with time, in log-scale, while the radius increases exponentially as well. The [G] profile at the beginning of the simulation (time *t*=0.25*d*, red trace in Fig. 9b), sees about half of the radius (corresponding to 7/8 of the total volume) above [G]_*V*_; however, during the exponential growth all cells have [G] way above the threshold concentration (time *t*=4*d*, black trace in the figure) with a nearly diffusive profile, a behavior that moreover does not depend on the value of the diffusion constant *D_G_* at long times. The [O] profile in this exponential growth regime is rather different from the previous example: as shown in Fig. 9c, it remains purely diffusional in shape (that is, similar to the results of Fig. 7 above), at all times. The radial distribution of the [ATP]/[ADP] ratio develops from an initial profile closely following the [G] distribution (red trace in Fig. 9d) into a flat distribution, with a much more peaked overshoot to high values close to the spheroid surface, compared to the case of surface-limited growth.

**Fig. 9.**
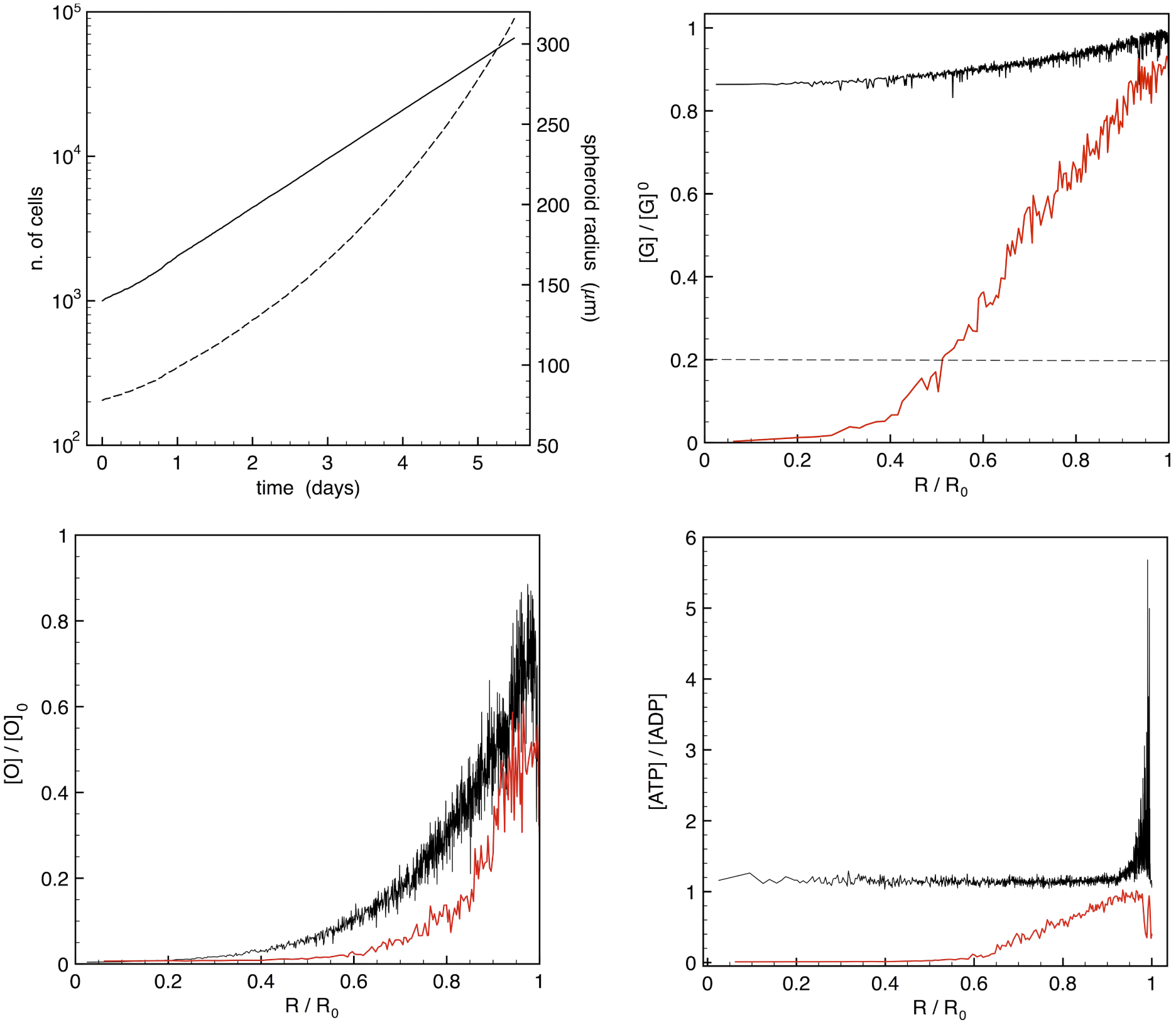
Simulation of 3D spheroid growth in the exponential regime. (a) Growth rate of the cell number (full, left ordinates) and of the spheroid radius (dashed, right ordinates), as a function of time. The linear rate of the cell number in log scale signifies exponential growth. (b) Glucose concentration ratio (units of the baseline concentration [G]_0_=25 mM) at times *t*=0.25 days (red) and 4 days (black), as a function of spheroid radius (normalized to the respective *R_max_* for each data set). The horizontal dashed line indicates the glucose viability threshold, [G]_*V*_, here equal to 0.2 [G]_0_. (c) Same as (b), for the oxygen ratio (with [O]_0_=0.6 mM). (d) Same as (b), for the [ATP]/[ADP] ratio. (Color online)

Taken together, the results of these two examples show that the 3D ABM retains the variety of responses that can be obtained with the simplified model, moreover with the added flexibility of active feedbacks linking the inter-cellular response to intra-cellular metabolite evolution. Notably, the examples above were limited to relatively small spheroids, containing up to about 60,000 cells. At such sizes, either the surface-limited, or the rapid exponential growth is observed in real spheroids. However, at larger sizes nearly all tumorsphere assay experiments demonstrate a saturation and eventually arrested growth [29, 30, 44], a behavior that is customarily described in simpler mathematical models by a capacity-limited growth function (Gompertzian or logistic). In the next Section 3.2 we will introduce additional features in the model, to account for the experimental observation of saturation at increasing sizes.

### 3.3 Simulation of spheroid growth saturation

The expansion of tumorspheres beyond a radius of 3-400 *μ*m brings about a number of changes in the cell response [29,30,44]. At this size, the central core first shows traces of necrosis. The size of the necrotic zone increases, until close to the arrest it would cover a large fraction of the total volume. Already a small fraction of necrotic core induces a reduction in glucose and oxygen utilization rates; in parallel, the average cell size may be reduced, the cell radius decreasing by as much as 5-10% of the initial value; at the same time, as the spheroids grow, the outer rim of proliferating cells shows a slight decrease in thickness [30]. It may be noticed that in these experiments it is usually difficult to separate between quiescent (G0) and proliferating cells, therefore the rim thickness is determined simply by difference with the size of the necrotic core, which in turn can be clearly observed e.g. by staining techniques [45]. A relevant observation from Freyer’s work is that spheroid size at equilibrium appears to be correlated with the size of the spheroid at the onset of necrosis.

Several tumor growth models have already proposed to include the release of a “necrosis factor” from the inner cells (see [46] and references therein). However, the phenomena relative to growth arrest seem to occur starting from the spheroid surface [29, 30], namely the drop in glucose and oxygen consumption and reduction in cell size, which altogether point to a modification of the cell homeostasis, once the necrosis starts somewhere in the volume. Such an effect could not be accounted only by the diffusion of such a “necrotic factor”, which necessarily would start from the inside of the sphere volume. Therefore, a feedback action involving some kind of cell-cell signaling (e.g., some “growth inhibitor” released from viable rim cells, such as lactate or other pH modifiers [47]) should be implicated in the process of arrested growth.

In the simulations reported in this last Section, we introduce two key elements in the ABM. The first element is the explicit distinction between necrotic, quiescent and viable cells. All definitions are based on the local levels of oxygen and glucose, implying respectively the stop (necrosis) or a reduction (quiescence) in the glycolysis, and consequently in the regeneration of ATP. The second element is the experimentally observed shedding of cells from the surface of tumor spheroids [5, 48, 49], similar to what happens in a real tumoral mass disseminating circulating cancer cells [50].

There are several methods to induce a quiescent (G_0_) cell status in experimental cultures, such as nutrient starvation for several days, culture to confluence, use of kinase inhibitors, each with different mechanisms by which they inhibit cell growth [51,52]. In our ABM model, the distinction among cell status is introduced based on probabilistic criteria, leading to Boolean variables to be set to ‘on’ or ‘off’. Tolerance times *t_C_*, *t_O_* and *t_G_* are introduced, by counting the time each cell is, respectively, found confluent (i.e., surrounded by at least two filled shells of neighbors), in low oxygen, or low glucose conditions. If a cell remains under such conditions for a continuous duration of time *t* larger than at least two combined tolerance times, it is switched to G0; moreover, if a cell remains continuously under low oxygen for a time *t >* 2*t_O_*, it becomes necrotic. The concentration thresholds for oxygen and glucose viability are fixed at the already cited values [O]_*V*_ =0.3 mM and [G]_*V*_ =5 mM. For *t_O_* and *t_G_*, we independently tested a range of values comprised between a few hours and 2 days. Figure 10 shows a simulation over 24 days of growth, for a spheroid starting with just 10 cells; the parameters allow to attain a doubling time-scale comparable to experimental observations, e.g. in EMT6 cells [29, 30]. From the upper panel, showing the number of cells, it can be seen that the growth starts with exponential regime, then starts saturating around *t*=5-6 days; however, at a later time (*t* ≃12 days) a second exponential regime takes over. This points at the fact that inclusion of cell differentiation (normal, quiescent, necrotic) is a necessary but not sufficient condition to induce saturation of the growth. Necrosis sets is some time after the initial growth, around *t*=3 days (dashed curves), and it is likely responsible for the initial apparent saturation. By looking at the volume growth rate (radius of the spheroid, shown in the lower panel) it is seen that the difference between outer and necrotic core becomes constant, i.e. the radius of the active cell rim is constant. This observation is in agreement with experimental observations [29, 30], although with the present set of parameters the size of the active rim is in the range 40-50 *μ*m, against experimental values about 2-4 times as large, for similar growth conditions (low-oxygen and [G] 2-20 mM) and doubling times.

**Fig. 10.**
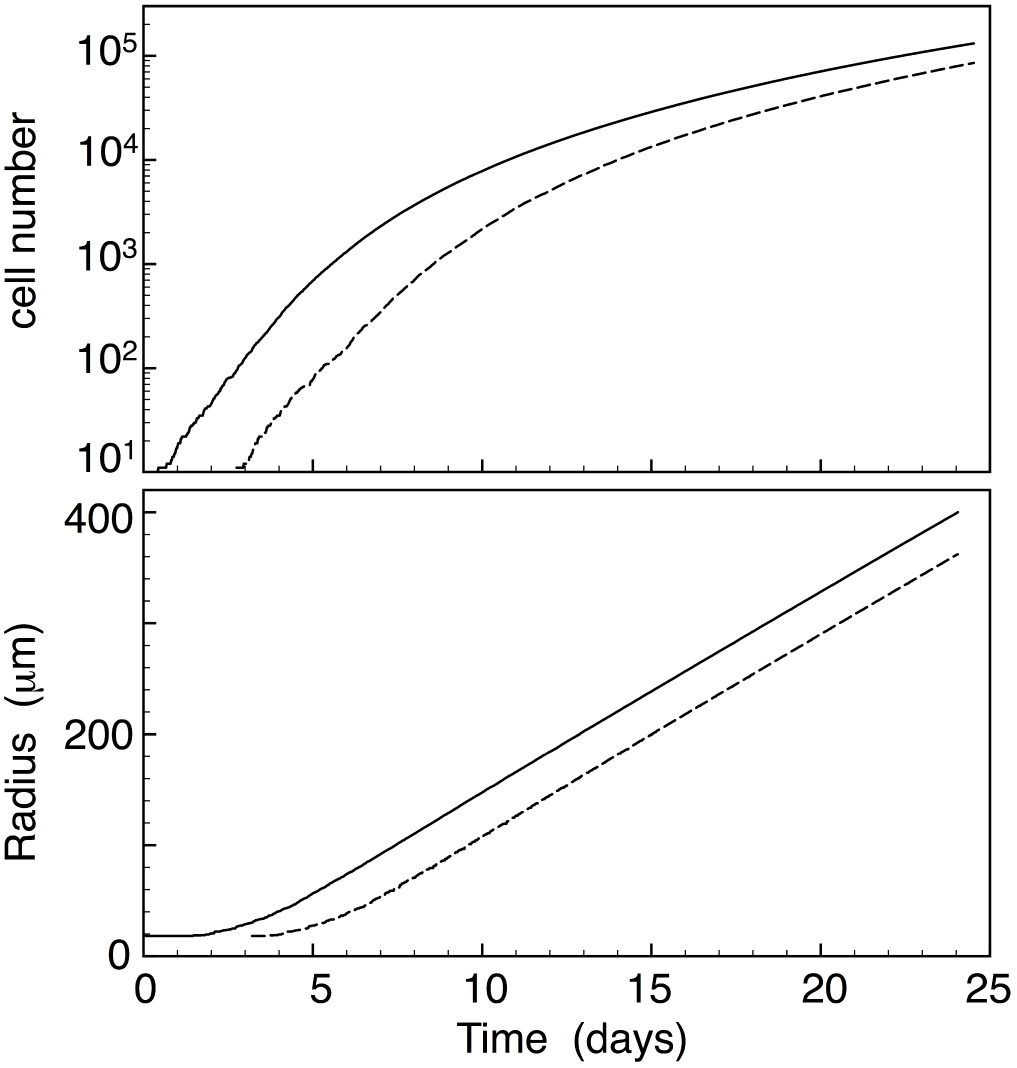
Simulation of long-term 3D spheroid growth including cell differentiation. Above: Total cell number (continuous) and number of cells in the necrotic core (dashed). Below: radius of the spheroid (continuous) and of the necrotic core (dashed).

Cell shedding from the surface of spheroids has been already included in some growth models [11, 12, 29, 53], as it has been considered a crucial element to attain equilibrium conditions. We include this possibility in the form of a continuous probability function depending on the discrete variable 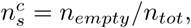, that is the ratio between the number of unoccupied and total neighbour sites of each cell *c*. The probability function has the simple form:

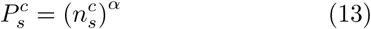

In this way, the shedding rate automatically depends on the spheroid surface (since the number of surface cells having *n_empty_ < n_tot_* is itself proportional to the surface), via the empirically adjustable parameter *α*. Moreover, as well according to experimental indications, shedding was started only after the spheroid attains a minimum size *R_s_*.

In Figure 11 we show the results of simulation runs using the same parameters as in Fig. 10, with and without cell shedding, with different values of *α*=8,10,12, and *R_s_*=50,100,200 *μ*m. The data points are from two different experiments on EMT6/Ro mouse mammary tumor cells [12]. After the initial exponential regime, a beginning of growth saturation effect is observed, leading to a sublinear growth rate. A correlation between the two parameters is observed, namely the larger *R_s_*, the larger is the smallest *α* for which growth starts saturating. The closest approximation to the experimental data is obtained for *R_s_*=125 *μ*m and *α* = 10.3. However, we also notice that growth saturation it is extremely sensitive to the numerical parameters, even a few percent variation of *α* changing the saturation behavior. Biological processes are instead robust and tolerant with respect to fluctuations, therefore such a saturation may rather arise as an artifact of the mathematical model; more effective mechanisms should be at work in the real spheroids, and be included in future developments of the model, such as the above cited controlled release of “growth inhibitors” by the cells.

**Fig. 11.**
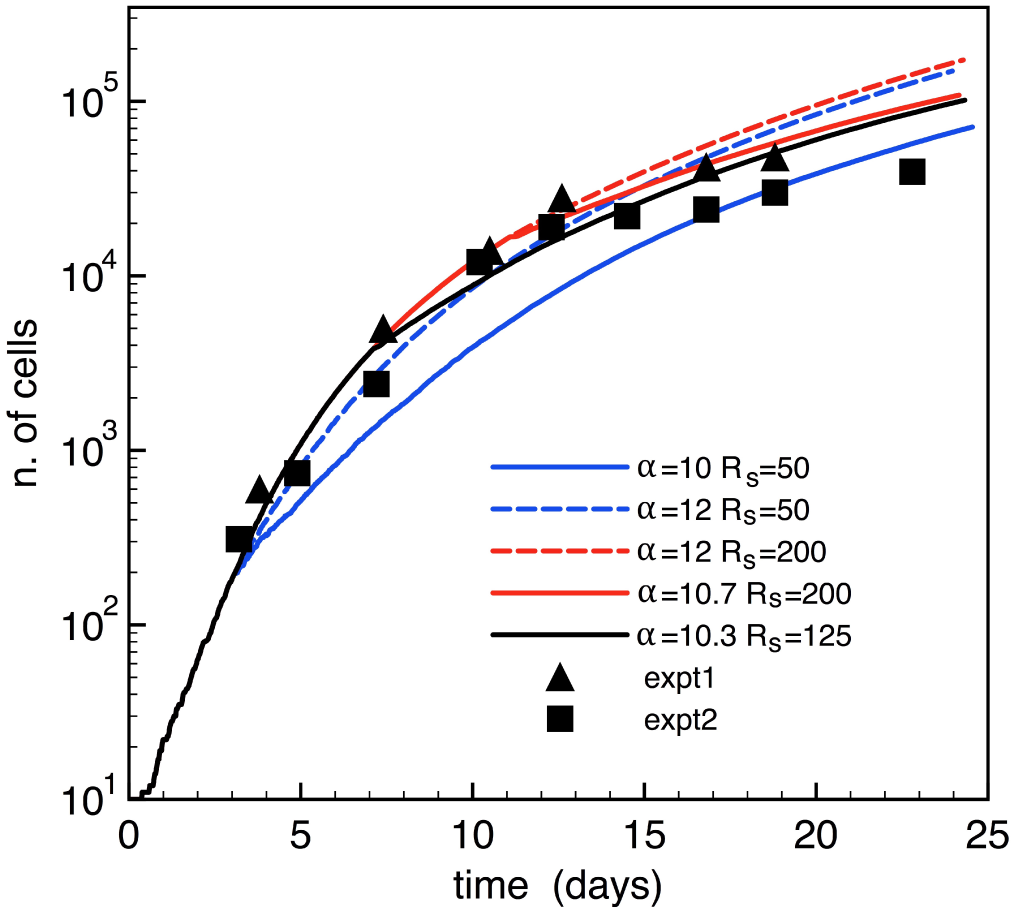
Simulation of long-term 3D spheroid growth including cell differentiation and cell shedding from the outer surface. Simulation parameters as given in the legend; experimental data from [12]. (Color online)

A further possibility that could be considered adding to the model is the role of mechanical stress originating from the external matrix, which is reported to be among the possible factors inhibiting growth when the matrix is made increasingly rigid, e.g. by controlled addition of agarose gel [54].

## 4 Conclusions

A multi-scale agent-based model (ABM) combined with a description of relevant signalling pathways has the potential of addressing basic research questions, and possibly identifying novel molecular targets [40, 55]. Mathematical modeling of the effects of biomedical treatments on tumor-sphere growth, such as chemo- and radiotheraphy, could be a game changer for the development of new, efficient therapies. In this work we extended to fully-3D the capabilities of our multicellular evolution ABM [24], with the aim of supporting the analysis of experiments of artificial growth of multicellular cancer spheroids, or “tumor-spheres”. The most important qualitative advance of the model is the inclusion of a local metabolic network, which allows each cell to independently regulate its ATP/ADP levels in response to the glucose/oxygen supply from the exterior, and to the ATP utilization rate for the cellular functions.

By filling a random 3D lattice of irregular polyhedra, the current model can mimic the growing tumorsphere, including cells with different phenotypes. Discrete variables with Boolean logic represent the switching off/on of specific “genes”, leading to probabilistic outcomes, empirically representing the de/activation of a given protein pathway. In our approach, the space- and time-dependent concentrations of the biochemical constituents, and gene activation/repression factors are computed numerically by non-linear partial differential equations (PDEs) describing the linked evolution of the various factors, such as the glycolysis/oxygen/ATP cycle. Overall, the model was designed so as to work with a relatively small number of adjustable parameters. The objective is to produce a working model especially aimed at guiding and interpreting selective experiments of stem-cell spheroid growth. At this stage, we did not yet attempt a strict comparison with experimental data on real cell lines. However, already at this level, some qualitative experiment/simulation comparison is demonstrated, based on empirical adjustment of the various parameters and rate constants. The results on both short-term and long-term growth of multicellular spheroids are very promising, for the future applications of the model to realistic conditions of biological relevance.

## Acknowledgements

I am grateful to Dr. C. Lagadec (INSERM Lille) for introducing me to the subjects of tumorspheres, cancer stem cells, and reprogramming. This work was developed thanks to partial funding from the SIRIC OncoLille under project “ModCel”. Computer resources provided by the CINES and IDRIS French Supercomputing Centres, under grant A0040707225.

